# Reduced Backward Alpha Propagation at Rest Marks the Autism Continuum

**DOI:** 10.64898/2026.05.09.723982

**Authors:** Luca Tarasi, Andrea Alamia, Vincenzo Romei

## Abstract

Autism spectrum disorder and subclinical variation along the autism continuum are characterized by in sensory processing and cognitive integration, phenomena increasingly linked to atypical large-scale communication across cortical hierarchies. While structural and functional connectivity differences have been extensively documented, whether autistic traits are associated with a reorganization of the directional properties of ongoing cortical activity remains less understood. Here, we recorded resting-state EEG from 201 young adults selected from the lower and upper terciles of the Autism Quotient distribution and analyzed traveling-wave dynamics over parieto-frontal lines. Individuals with higher autistic traits showed a selective shift in left-hemisphere alpha-band traveling-wave directionality, driven primarily by reduced backward-dominant propagation and accompanied by a reciprocal shift toward forward dominance. This effect was anatomically specific, absent in an occipito-central control line set, and not accompanied by a matching pattern of group differences in oscillatory power, aperiodic spectral parameters, or peak alpha frequency. These findings identify resting-state alpha traveling waves as a candidate physiological signature of altered directional organization across the autism continuum, with potential relevance as a trait-sensitive neural marker.

## Introduction

Autism spectrum disorder (ASD) is a neurodevelopmental condition characterized by differences in social communication, restricted or repetitive behaviors and interests, and atypical sensory processing (APA, 2013). Beyond these behavioral features, a central open question concerns how variation across the autism spectrum relates to the organization of large-scale brain communication^1,2^. Although a substantial literature has examined atypical structural and functional connectivity in autism^3–5^ much less is known about whether autistic traits are associated not only with changes in the strength of large-scale interactions, but also with changes in their directional organization across cortical hierarchies. Here, we address this question from a dimensional perspective, focusing on autistic traits in a non-clinical sample.

One influential framework for interpreting autistic features is predictive processing, according to which perception and action emerge from the continuous interaction between prior expectations and incoming sensory input^6,7^. Within this broad framework, several accounts have proposed that autistic perception may reflect an atypical balance between prior beliefs and sensory input^8,9^, whether described in terms of weakened priors^10^ or altered precision weighting^11^. These accounts are formulated at a computational level, but they do not directly specify how such an imbalance should appear in the directional dynamics of large-scale cortical activity. Addressing this gap requires a neural measure that captures not only whether distant regions interact, but also the direction in which rhythmic activity propagates across the cortical hierarchy.

Rhythm-based models of cortical communication provide a useful bridge between these computational proposals and measurable neural dynamics. In these models, alpha-band activity has often been linked to feedback or top-down signaling, whereas higher-frequency activity has been associated with feedforward transmission of sensory evidence and prediction errors (Figure 1A)^12–16^. Although alpha rhythms are functionally heterogeneous and cannot be reduced to a single role^17–25^, their large-scale directional component is especially relevant here because it has been proposed to reflect feedback-like communication along cortical hierarchies^15,26,27^. If autistic traits are associated with a relative attenuation of such feedback-like signaling^28–31^, then this should be expressed as a shift in the directional balance of ongoing rhythmic propagation, potentially favoring forward over backward traveling activity (Figure 1B)^32^.

**Figure 1.**
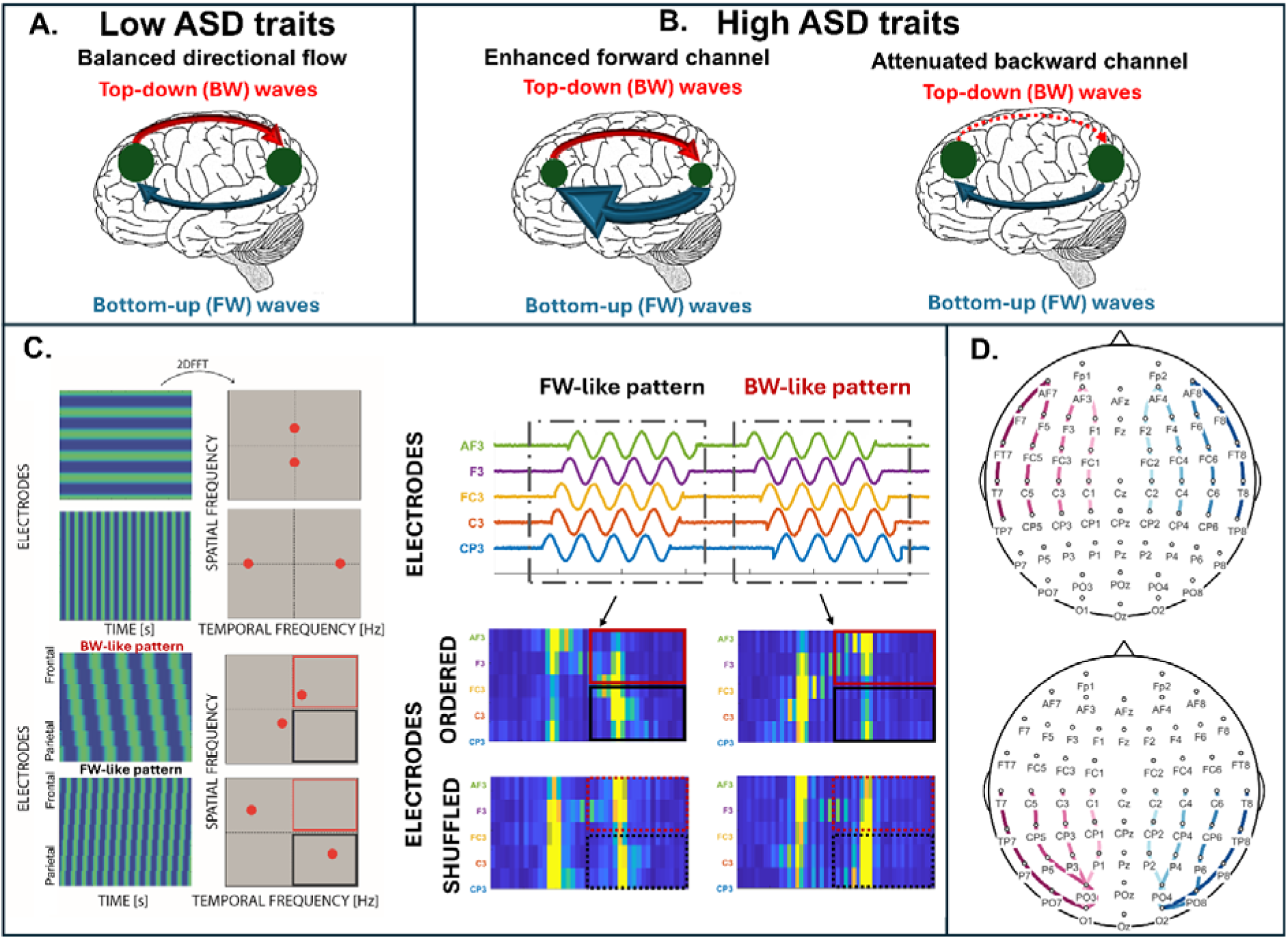
Hypothesis, methodological framework, and electrode montage. (A) Two architectural scenarios for the directional reorganization of traveling waves in autistic traits. In individuals with low autistic traits (left), backward (top-down, red) and forward (bottom-up, blue) traveling waves over the parieto-frontal network are organized in a balanced directional regime. In individuals with high autistic traits, two alternative scenarios could in principle produce an altered directional balance: an enhanced forward channel (centre), in which forward propagation is amplified and increases the relative weight of bottom-up signals, or an attenuated backward channel (right), in which backward propagation is weakened, with the forward component shifting reciprocally as a consequence of the tightly coupled push–pull regime in which the two directions operate. The attenuated backward channel represents the a priori scenario under predictive-processing accounts of autism. However, the two scenarios are not mutually exclusive but make distinct predictions about which propagation direction primarily carries the effect; as detailed in the Results, the present data most strongly support the attenuated backward channel scenario, with the forward shift appearing weaker and likely reciprocal. (B) Detection of traveling waves via two-dimensional Fourier transform (2D-FFT). Applying a 2D-FFT to a space–time matrix of EEG activity (electrodes along one axis, time along the other) produces a spectral representation in which the position of spectral peaks reflects both the temporal frequency of the oscillation and the direction of its propagation across the electrode line. Stationary waves (top) yield peaks aligned along the spatial-frequency axis, whereas waves propagating along the line (bottom) shift the peaks into one of the two off-diagonal quadrants depending on their direction of propagation. The upper-left quadrant (red) captures backward (BW)-like patterns travelling from posterior to anterior sites, while the lower-right quadrant (black) captures forward (FW)-like patterns travelling from anterior to posterior sites. (C) Application to EEG data and surrogate control. Single-trial EEG segments from a representative parieto-frontal line are first arranged in their natural anatomical order (“Ordered” rows), and the 2D-FFT is computed within each segment. Examples of FW-like and BW-like patterns are shown together with their corresponding spectral signatures, in which the relevant quadrant is preferentially populated. To control for non-directional spectral features (e.g., 1/f background, broadband power), the same procedure is repeated after randomly shuffling the electrode order along the spatial axis (“Shuffled” rows): in this case, any genuine directional signal is destroyed, and the residual spectral content reflects non-directional spectral structure rather than consistent propagation direction. (D) Electrode montage. Top: the eight parieto-frontal lines analyzed in the main analyses, four per hemisphere, each composed of five electrodes spanning the antero-posterior axis from temporo-parietal to fronto-anterior sites (left hemisphere in warm magenta tones, right hemisphere in blue tones). Bottom: the eight occipito-central control lines, used to test whether the effects observed over the parieto-frontal lines generalized to a distinct posterior control territory. Electrode labels and full line definitions are reported in the Methods.

Traveling-wave analyses are well-suited to test this idea. Rather than treating oscillatory activity as locally stationary, they quantify spatiotemporal patterns that propagate across cortical territory and dissociate forward and backward components within the same frequency band^33–37^. This is an important advantage over local power estimates or directionally agnostic connectivity measures, which do not directly isolate propagation direction. We focused on resting-state EEG because spontaneous activity provides a readout of intrinsic network organization in the absence of specific task demands^38–41^. If autistic traits are associated with a stable bias in the balance between feedback-like and feedforward-like signaling, this bias may already be detectable at rest as a directional predisposition of the system.

In the present study, we analyzed resting-state EEG traveling waves in a large sample of young adults selected from the lower and upper terciles of the Autism Quotient distribution, a strategy adopted to maximize statistical contrast in a sub-clinical trait distribution that is typically skewed toward low values^42^. We focused on posterior-to-frontal propagation within a parieto-frontal configuration, given the proposed role of parieto-frontal systems in attention, cognitive control, and the integration of internally generated models with ongoing processing^43–45^. We therefore focused primarily on the possibility of reduced backward-dominant alpha propagation, compatible with attenuated feedback-like signaling, while also considering the alternative possibility that directional imbalance could arise from a primary enhancement of forward propagation (Figure 1B). These possibilities are not mutually exclusive, and traveling-wave decomposition allows their relative contribution to be estimated rather than collapsed into a generic notion of imbalance^17,34^. We further asked whether any directional differences would be anatomically specific or broadly distributed across the scalp, and whether they would be accompanied by group differences in conventional spectral descriptors such as band-limited power, aperiodic signal properties, or individual alpha peak frequency. To test this directly, we compared a parieto-frontal line set with a posterior occipito-central control line set matched in structure but sampling a distinct scalp territory. The results revealed that higher autistic traits are associated with a reorganization of parieto-frontal alpha traveling waves, primarily driven by attenuated backward-dominant propagation, with the accompanying shift in forward propagation reflecting the strong reciprocal coupling between the two directions. This reorganization was anatomically specific, absent in an occipito-central control line set, and was not accompanied by a parallel pattern of group differences in band-limited power, aperiodic spectral parameters, or individual alpha frequency.

## Results

### Participants and design overview

To test whether resting-state traveling waves over a parieto-frontal line set carry a signature of autistic traits, we recorded EEG from 201 participants selected from a larger sample of 323 young adults on the basis of their Autism Quotient (AQ^46^) Social Interaction scores^47^. Participants were divided into two groups corresponding to the lower and upper terciles of the AQ distribution: a Low-AQ group (N = 95, mean AQ = 6.67 ± 0.20; mean age = 22.87 ± 0.30 years) and a High-AQ group (N = 106, mean AQ = 19.82 ± 0.37; mean age = 23.39 ± 0.30 years). The two groups did not differ in age (*t*(199) = 0.96, *p* = .338, BF_10_= 0.24) or in sex distribution (Low-AQ: 68 Female; High-AQ: 63 Female; X2(1) = 3.26, *p* = .071, BF_10_= 0.92).

### Left-alpha traveling-wave directionality differs between AQ groups

We first asked whether group differences in traveling-wave directionality were broadly distributed across frequencies, hemispheres, and line positions, or instead confined to specific spectro-topographic compartments (Figure 1C–D). The omnibus linear mixed-effects model on the parieto-frontal directionality index (*DI*; defined as the difference between band-averaged FW and BW wave amplitudes after surrogate normalization, such that positive values indicate relatively greater forward-wave dominance, negative values relatively greater backward-wave dominance, and values near zero a more balanced profile; Table S1) supported the latter possibility. DI varied across frequency bands (F(3, 6169) = 113.29, p < .001), hemispheres (F(1, 6169) = 14.86, p < .001), and medio-lateral line positions (F(3, 6169) = 51.98, p < .001), indicating that directional balance depended on both spectral and spatial factors. Crucially, the model revealed a Group × Band interaction (F(3, 6169) = 7.59, p < .001) and a Group × Band × Hemisphere interaction (F(3, 6169) = 3.64, p = .012), indicating that the association between AQ group and directional balance differed across frequencies and hemispheres. By contrast, the main effect of Group was not significant (F(1, 199) = 3.41, p = .066), arguing against a diffuse shift of *DI* across the parieto-frontal set. Importantly, the Group × Band × Hemisphere × Line Position interaction was null (F(9, 6169) = 0.62, p = .782), indicating that the critical spectro-hemispheric pattern did not further depend on medio-lateral line position.

To unpack this three-way interaction, we therefore adopted a hierarchical follow-up strategy. Because the four-way interaction involving Line Position was not significant, *DI* values were first averaged across line positions within subject, and the Group × Band interaction was then tested separately within each hemisphere (Table S2). This interaction was significant in the left hemisphere (F(3, 597) = 4.15, p = .006), but not in the right hemisphere (F(3, 597) = 0.69, p = .558), localizing the omnibus effect to the left parieto-frontal lines. We then followed up the left-hemisphere model with band-specific between-group contrasts, correcting across bands with Holm–Bonferroni (Table S3). A significant group difference emerged in the left-hemisphere alpha band, where low-AQ participants showed greater backward dominance than high-AQ participants (i.e., lower DI values: DI_LOW = −0.053, DI_HIGH = +0.020; t(199) = −2.73, p = .007, Holm-corrected p = .028, d = −0.38, BF = 4.77). No group differences were observed in the other left-hemisphere bands (theta: t(199) = −0.32, p = .750, BF10 = 0.16; beta: t(199) = −0.49, p = .622, BF10 = 0.17; gamma: t(199) = 0.09, p = .928, BF10 = 0.15; all Holm-corrected p = 1), with Bayesian evidence consistently favoring the null. Thus, the group effect on directional balance was not broadly distributed across the spectrum, but was selectively concentrated in the alpha band of the left hemisphere. As a robustness check, we recomputed directional balance using a direct FW/BW metric without surrogate normalization (Table S4–S6). All critical effects were preserved, including the alpha-left simple contrast (*t*(199) = −2.64, Holm-corrected *p*= .036, BF_10_= 3.88).

### FW/BW decomposition indicates that the left-alpha effect is driven primarily by reduced backward propagation

The group difference in DI could reflect a change in forward-wave amplitude, a change in backward-wave amplitude, or a shift in their relative balance. To distinguish among these possibilities, we decomposed the left-hemisphere alpha effect into forward (FW) and backward (BW) components using a repeated-measures ANOVA with Direction as a within-subject factor and Group as a between-subject factor (Figure 2C–D).

**Figure 2.**
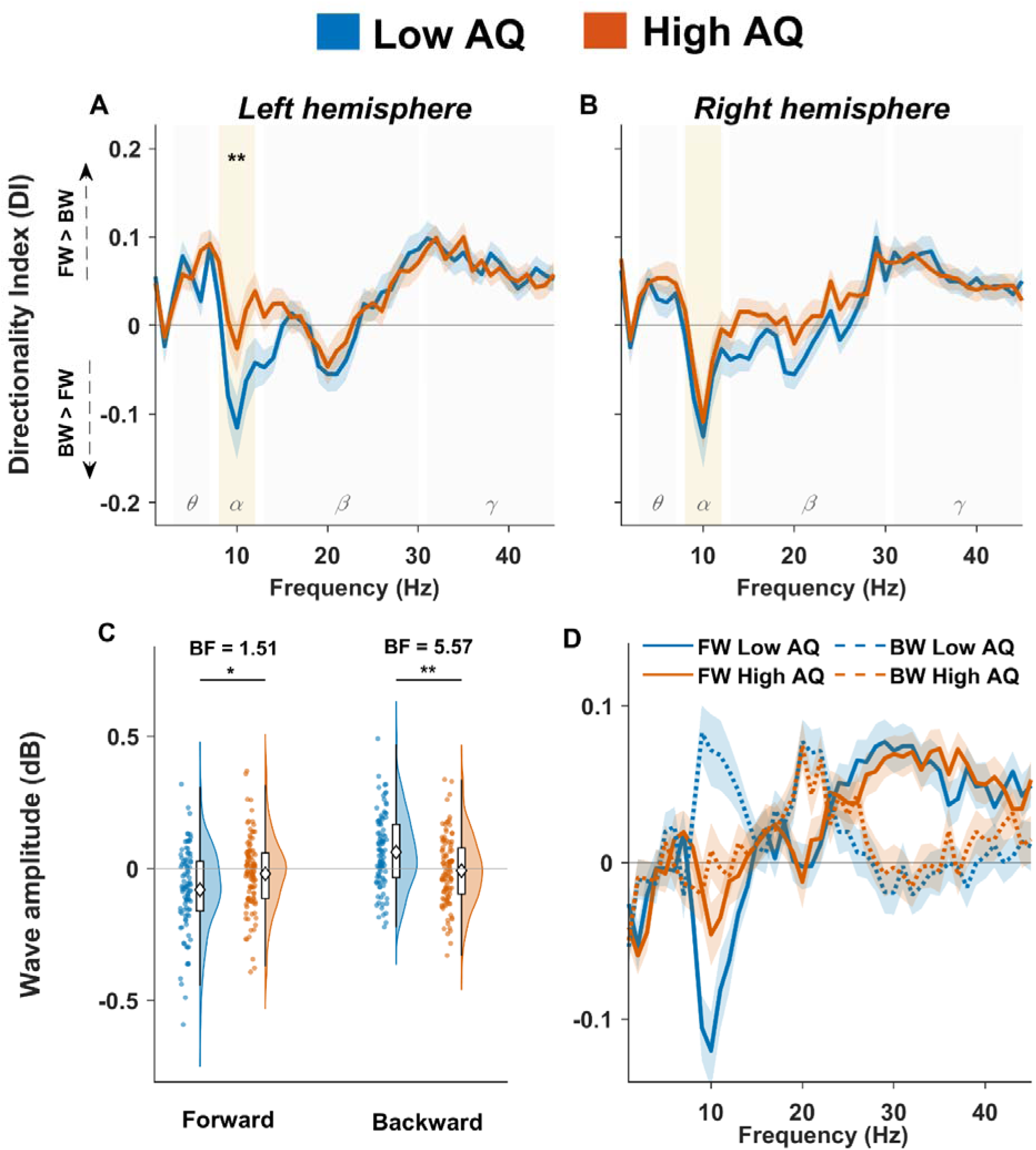
Parieto-frontal traveling-wave directionality differs between AQ groups in the left alpha band. **(A, B)** Directionality index (DI; computed as the difference between FW and BW wave amplitudes after surrogate normalization, in dB) as a function of frequency, averaged across the four parieto-frontal lines of the left (A) and right (B) hemisphere. Solid lines show group means for low-AQ (blue) and high-AQ (orange) participants; shaded areas denote ± 1 SEM. Canonical frequency bands (θ, α, β, γ) are indicated by background shading, with the alpha band highlighted. Positive DI values reflect forward-dominant propagation, negative values backward-dominant propagation. In the left hemisphere, the two groups diverge selectively in the alpha range, with low-AQ participants showing a shift toward backward dominance relative to high-AQ participants. The asterisk marks the alpha-left contrast surviving Holm correction in the hierarchical follow-up analysis. No comparable group difference is observed in the right hemisphere. **(C)** Decomposition of the alpha-left effect into FW and BW components. Raincloud plots show the distribution of single-subject FW and BW amplitudes (at the single left-hemisphere line showing the largest group effect on alpha-band DI), combining a half-violin kernel density estimate, jittered individual data points, a compact boxplot (median, interquartile range, whiskers at 1.5 × IQR), and the group mean (white diamond). Statistics reported in the text were computed after averaging across left-hemisphere line positions; the panel is intended to illustrate the asymmetric directional rebalancing underlying the alpha-left DI effect. High-AQ participants show reduced backward waves together with a weaker reciprocal shift in forward waves, consistent with the stronger Bayesian evidence for the BW contrast than for the FW contrast. Significance asterisks report two-tailed p-values from independent-samples t-tests. **(D)** Frequency-resolved forward and backward wave spectra at the single left-hemisphere line showing the largest group effect on alpha-band DI, shown for visualization purposes. Solid lines indicate forward waves, dashed lines backward waves; colors code AQ group as in panels A–C. Shaded areas denote ± 1 SEM. The alpha band is highlighted. The reorganization summarized in panel C is localized within the alpha range and takes the form of a crossover: low-AQ participants exhibit weaker forward and stronger backward propagation, whereas high-AQ participants show the opposite pattern, with the two spectra converging or reversing within the alpha band.

For the left-hemisphere alpha band, the Group × Direction interaction was significant (F(1, 199) = 7.43, p = .007, partial η^2^ = .036), indicating that the two groups differed in the relative balance between forward and backward propagation rather than in overall wave amplitude. Follow-up tests showed that BW alpha was reduced in the high-AQ group relative to the low-AQ group (LOW = +0.021, HIGH = −0.018; t(199) = 2.79, p = .006, d = 0.39, 95% bootstrap CI [0.10, 0.70], BF10 = 5.57). FW alpha showed a reciprocal shift in the opposite direction, being slightly larger in the high-AQ group than in the low-AQ group (LOW = −0.032, HIGH = +0.002; t(199) = −2.22, p = .028, d = −0.31, 95% bootstrap CI [−0.62, −0.04]), but Bayesian evidence for this contrast remained only anecdotal (BF10 = 1.51). The asymmetric Bayesian evidence is informative: support for a reduction in BW alpha was moderate (BF10 = 5.57), whereas support for the reciprocal FW increase remained anecdotal (BF10 = 1.51), suggesting that the primary locus of the effect lies in the backward component.

Across participants, FW and BW alpha values in the left hemisphere were strongly anticorrelated (r = −0.682, p < .001), consistent with a reciprocal push-pull relationship between the two directions. When examined separately by group, this anticorrelation was stronger in low-AQ participants (r = −0.747) than in high-AQ participants (r = −0.585, z = −2.06, p = 0.039). We therefore interpret the correlation analysis as descriptive support for a reciprocal directional regime, with stronger reciprocal coupling in the low-AQ than in the high-AQ group.

### Group differences in traveling-wave directionality were not explained by spectral power, aperiodic parameters, or peak alpha frequency

We next examined whether the group difference in traveling-wave directionality was accompanied by broader group differences in conventional spectral features derived from the same parieto-frontal line spectra used in the TW analysis, namely band-limited power, aperiodic spectral parameters, and peak alpha frequency (PAF) (Figure 3). For band-limited power, we used the same factorial structure as in the main *DI* analysis, estimating power separately for each parieto-frontal line and testing Group, Band, Hemisphere, and Line Position effects in a linear mixed-effects model (Table S7). The model recovered the expected effects of frequency band (F(3, 6169) = 15179, p < .001) and line position (F(3, 6169) = 495.05, p < .001), together with a Band × Line Position interaction (F(9, 6169) = 71.44, p < .001), indicating that oscillatory amplitude varied across the parieto-frontal set as a function of frequency and medio-lateral position. A Group × Band interaction was also present (F(3, 6169) = 3.23, p = .022), but no higher-order interaction involving Group approached significance (all F < 1.31, all p > .27), including the critical Group × Band × Hemisphere term (F(3, 6169) = 0.07, p = .974). Follow-up contrasts on power averaged across hemispheres and line positions did not reveal reliable group differences in any band after Holm correction (Table S8; theta: t(199) = 1.50, p = .135, BF10 = 0.44; alpha: t(199) = 1.60, p = .111, BF10 = 0.51; beta: t(199) = 1.75, p = .081, BF10 = 0.64; gamma: t(199) = 1.40, p = .164, BF10 = 0.38; all Holm-corrected p > .32). Thus, the two AQ groups showed broadly comparable oscillatory-amplitude profiles in the parieto-frontal line spectra.

**Figure 3.**
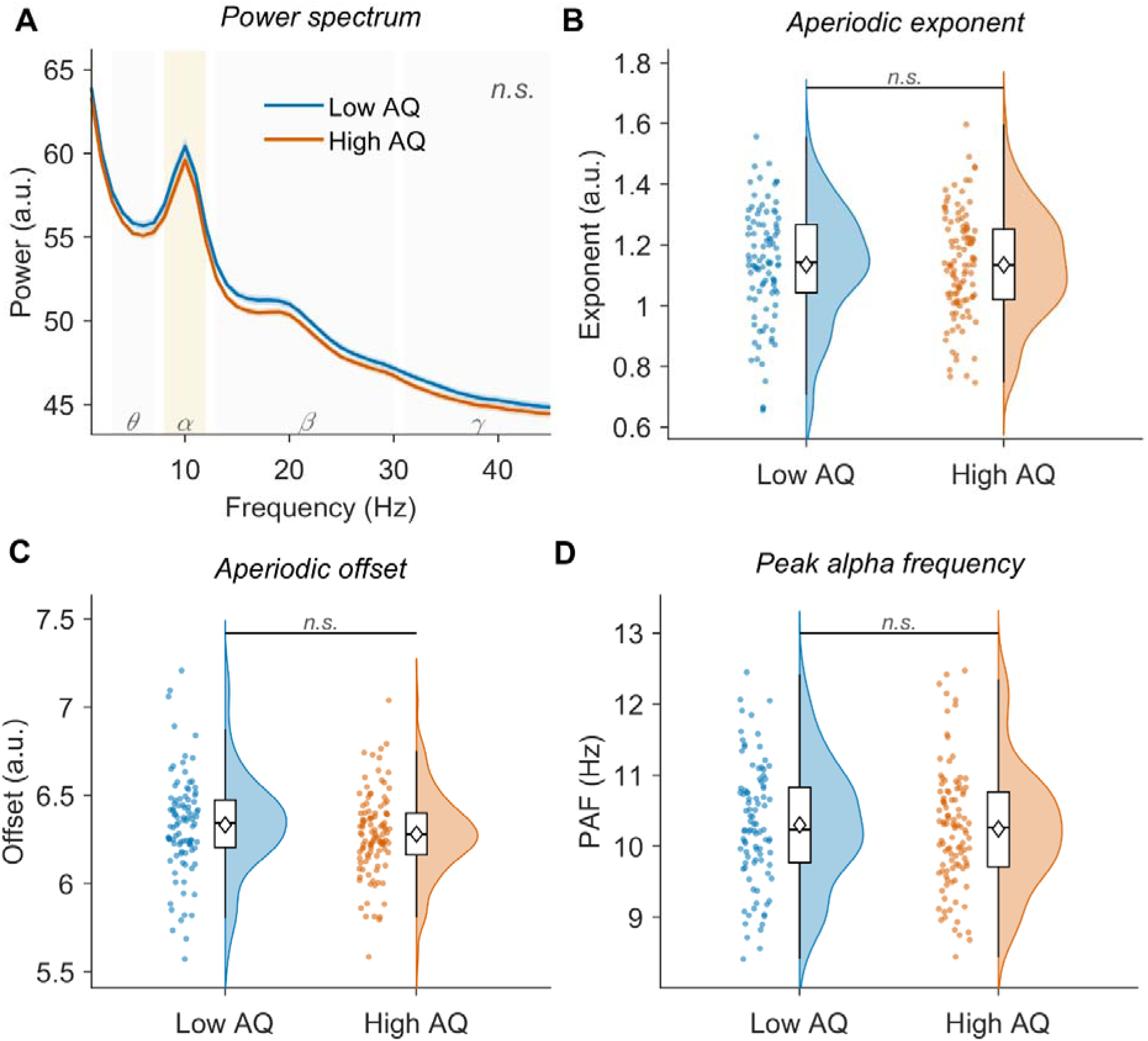
The parieto-frontal traveling-wave effect was not paralleled by conventional spectral measures. (A) Band-limited power spectra from the same parieto-frontal line set used in the traveling-wave analysis, shown separately for low-AQ (blue) and high-AQ (orange) participants. Solid lines indicate group means; shaded areas denote ± 1 SEM. Canonical frequency bands (θ, α, β, γ) are indicated by background shading, with the alpha band highlighted. Although the omnibus mixed-effects model showed the expected effects of band, no band-specific group contrast survived Holm correction (see Results and Table S7; S8). (B–D) Aperiodic and periodic spectral parameters derived from FOOOF fits to the same parieto-frontal line spectra used in the TW analysis. Panels show visualization summaries of the data, whereas inferential statistics were performed with line-wise mixed-effects models including Group, Hemisphere, and Line Position (see Results and Table S9; S10). (B) Aperiodic exponent. The slope of the 1/f background did not show a reliable main effect of Group or any interaction involving Group and Hemisphere; although exponent varied across line positions, no robust group-related pattern emerged. (C) Aperiodic offset. The vertical position of the 1/f background likewise showed no reliable effect involving Group. (D) Peak alpha frequency (PAF), defined as the centre frequency of the highest-power periodic component falling within 7–13 Hz after removal of the aperiodic background. PAF did not differ reliably between groups and did not show any interaction involving Group, although modest variation across line positions was present overall. Taken together, these complementary spectral controls indicate that the left-alpha traveling-wave effect was not accompanied by a parallel pattern of group differences in conventional spectral descriptors of the same parieto-frontal line spectra. Equivalent analyses performed on the occipito-central control line set likewise yielded no group-related pattern consistent with the main TW effect (Supplementary Figure S2).

We then used FOOOF ^48^ to parameterize the same parieto-frontal line spectra into aperiodic and periodic components (Table S9). Model fits were of high quality across the dataset (mean R² = 0.981, SD = 0.037; mean fit error = 0.046, SD = 0.021), with no failed fits. For the aperiodic exponent, there was no main effect of Group (F(1, 199) < 0.001, p = .983), no Group × Hemisphere interaction (F(1, 1393) = 0.04, p = .847), and no Group × Hemisphere × Line Position interaction (F(3, 1393) = 0.14, p = .933). A Group × Line Position interaction was observed (F(3, 1393) = 2.68, p = .046), but follow-up contrasts at each line position were all null after correction (Table S10; all Holm-corrected p = 1; all BF10 < 0.19), indicating no robust group difference at any individual line. For the aperiodic offset, no reliable effect involving Group was observed (Group: F(1, 199) = 2.03, p = .156; Group × Hemisphere: F(1, 1393) = 0.36, p = .550; Group × Line Position: F(3, 1393) = 1.08, p = .358; Group × Hemisphere × Line Position: F(3, 1393) = 0.52, p = .668).

Finally, we tested whether the left-alpha directionality effect was accompanied by a shift in peak alpha frequency (PAF), again estimated from the same parieto-frontal line spectra after FOOOF decomposition. PAF did not differ between groups (F(1, 194.95) = 0.20, p = .653), or any interaction involving Group (all p > .44). Taken together, these analyses indicate that the observed TW group effect was not accompanied by a parallel pattern of group differences in conventional spectral descriptors of the same parieto-frontal line spectra.

### The effect did not generalize to a distinct occipito-central control line set

To test whether the parieto-frontal effect reflected a spatially specific phenomenon rather than a diffuse scalp-wide pattern, we repeated the *DI* analysis in a distinct occipito-central control line set (Figure 1D, S1). This control set consisted of eight posteriorly shifted five-electrode lines, again arranged along the antero-posterior axis and matched in number and structure to the main parieto-frontal lines, but sampling a different scalp territory extending from occipital to central sites. The omnibus linear mixed-effects model on occipito-central *DI* showed no effect of Group (F(1, 199) = 0.02, p = .898) and no Group × Band (F(3, 6169) = 1.40, p = .242), Group × Hemisphere (F(1, 6169) = 0.01, p = .925), or Group × Band × Hemisphere interaction (F(3, 6169) = 0.13, p = .943) (Table S11). No interaction involving Group approached significance, including the Group × Band × Hemisphere × Line Position term (F(9, 6169) = 0.58, p = .818). Because no group-related term was significant, no hierarchical follow-up or FW/BW decomposition was performed. Thus, the group difference observed in the parieto-frontal line set did not generalize to this posterior control set. We next examined whether the occipito-central control set showed any broader group differences in conventional spectral features derived from the same line spectra. For band-limited power (Table S12), the mixed-effects model recovered the expected effects of frequency band (F(3, 6169) = 19939, p < .001), hemisphere (F(1, 6169) = 40.72, p < .001), and Line Position (F(3, 6169) = 274.01, p < .001), together with a Band × Line Position interaction (F(9, 6169) = 22.71, p < .001), indicating structured spectral variation across the occipito-central set. A Group × Band interaction was also present (F(3, 6169) = 3.52, p = .014), but no higher-order interaction involving Group approached significance, including Group × Band × Hemisphere (F(3, 6169) = 0.14, p = .934), Group × Band × Line Position (F(9, 6169) = 0.06, p = 1.000), and Group × Band × Hemisphere × Line Position (F(9, 6169) = 0.03, p = 1.000). Follow-up contrasts averaged across hemispheres and line positions did not reveal group differences in any band after Holm correction (Table S13; theta: t(199) = 1.53, p = .126, BF10 = 0.46; alpha: t(199) = 1.41, p = .161, BF10 = 0.39; beta: t(199) = 1.69, p = .092, BF10 = 0.59; gamma: t(199) = 1.27, p = .205, BF10 = 0.33; all Holm-corrected p > .36). We then used FOOOF to parameterize the same occipito-central line spectra into aperiodic and periodic components (Table S14). Model fits were of high quality across the dataset (mean R^2^ = 0.986, SD = 0.014; mean fit error = 0.051, SD = 0.021), with no failed fits. For the aperiodic exponent, there was no main effect of Group (F(1, 199) = 0.02, p = .884), no Group × Hemisphere interaction (F(1, 1393) = 1.91, p = .167), no Group × Line Position interaction (F(3, 1393) = 0.26, p = .851), and no Group × Hemisphere × Line Position interaction (F(3, 1393) = 0.10, p = .958). The same was true for the aperiodic offset (Group: F(1, 199) = 2.14, p = .146; Group × Hemisphere: F(1, 1393) = 0.82, p = .367; Group × Line Position : F(3, 1393) = 0.32, p = .812; Group × Hemisphere × Line Position : F(3, 1393) = 1.24, p = .293). Finally, we asked whether the control set showed any group-related shift in peak alpha frequency, again estimated line-wise after FOOOF decomposition. PAF did not differ between groups (F(1, 197.41) = 0.19, p = .663), hemispheres (F(1, 1368.6) = 2.10, p = .148), or any interaction involving Group (Group × Hemisphere: F(1, 1368.6) = 0.65, p = .420; Group × Line Position : F(3, 1368.5) = 0.37, p = .774; Group × Hemisphere × Line Position : F(3, 1368.5) = 0.94, p = .421), although PAF varied modestly across line positions overall (F(3, 1368.5) = 9.52, p < .001). Taken together, these analyses indicate that the null result in the occipito-central control set was not masked by an alternative pattern of group differences in conventional spectral descriptors.

## Discussion

The present study shows that autistic traits are associated with a selective reorganization of resting-state traveling-wave directionality over a parieto-frontal line set. The main effect was observed in the left-hemisphere alpha band, where individuals with higher autistic traits showed reduced backward-dominant propagation together with a reciprocal shift toward forward dominance. More specifically, the effect was confined to alpha-band directionality in the left parieto-frontal lines and was not observed either in other frequency bands or in the posterior occipito-central control lines. Moreover, the asymmetric weight of evidence across the two directions is informative: Bayesian analyses indicate that the group difference was driven primarily by the backward component, with forward propagation shifting in parallel within a strongly reciprocal directional regime.

This pattern is broadly consistent with previous findings in clinical ASD populations^49^. Adults with ASD have been reported to show reduced alpha synchronization to behaviourally relevant stimuli^50^, as well as reduced alpha-band connectivity during tasks engaging cognitive control, particularly in networks involving inferior frontal regions^51^. Seymour et al.^52^ likewise reported reduced alpha-band feedback connectivity during visual processing in ASD, with higher autistic traits associated with weaker feedback signaling. Importantly, recent work has extended this pattern to traveling-wave dynamics themselves: during a visual entrainment task, neurotypical adults showed an increase in backward waves, whereas adults with ASD exhibited the opposite profile, namely an increase in forward waves at the entrained frequency^53^. Because backward and forward traveling waves have been proposed to reflect predictive/feedback-like and sensory/feedforward-like signaling, respectively, that finding is consistent with a relative bias toward bottom-up propagation during stimulus-driven processing in ASD. Our results complement this task-based evidence by suggesting that a related directional asymmetry may already be detectable at rest, in the absence of external stimulation, and may be expressed not only in the strength of oscillatory coupling but also in the intrinsic directional organization of large-scale wave propagation.

A natural framework for interpreting these findings is predictive processing, according to which perception arises from the continuous interaction between prior expectations and incoming sensory evidence ^6,7,54^. Within rhythm-based formulations of this framework, alpha-band dynamics have often been linked to backward or feedback-like signaling, whereas higher-frequency activity has been associated with forward or feedforward-like transmission of sensory evidence and prediction errors^12,13,15,27,55,56^. From this perspective, the selective reduction of backward alpha propagation observed here is compatible with the idea that individuals with higher autistic traits differ in the balance between internally generated and sensory-driven processing, in line with accounts emphasizing attenuated priors or altered precision weighting^10,11,57–60^.

At the same time, the present data did not directly demonstrate altered prior weighting or predictive inference. Our measurements were obtained at rest, in the absence of any stimulus or task, and therefore cannot index how priors and prediction errors are dynamically weighted during active perception. What they do characterize is the intrinsic directional organization of ongoing alpha activity. In this sense, the present findings are better understood as evidence for a resting-state architectural bias that is compatible with reduced backward, feedback-like signaling, rather than as a direct neural signature of altered predictive computation itself. This interpretation is strengthened by the asymmetric evidential profile of the two propagation directions: the stronger evidence concerned reduced backward alpha, whereas the forward increase was weaker and plausibly secondary to the strong reciprocal coupling between the two components. Read in this way, the present results are more readily accommodated by an account emphasizing reduced backward/feedback-like signaling than by one positing a primary enhancement of forward/feedforward-like propagation. Consistent with this view, recent work from our group has shown that individuals with higher autistic traits exhibit a reduced ability to flexibly modulate alpha activity following explicit prior induction^61^, suggesting that the resting-state attenuation described here may translate into measurable differences when a task explicitly recruits backward-based processing^62^.

A second notable feature of the findings is their anatomical and spectral specificity. The effect was localized in the left parieto-frontal lines, where the high-AQ group showed reduced backward alpha together with a shift toward forward dominance. No comparable reorganization was observed in the posterior occipito-central control lines, arguing against a diffuse scalp-wide phenomenon and instead pointing to a circumscribed alteration in parieto-frontal directional dynamics. One plausible interpretation is that the parieto-frontal line set more directly samples large-scale posterior-to-anterior interactions and is therefore better positioned to capture individual differences in the balance between backward, feedback-like propagation and its reciprocal forward component. By contrast, the occipito-central control lines sample a more posterior scalp territory and may reflect a different balance of local sensory and shorter-range dynamics. At a functional level, this localization is compatible with the proposed role of parieto-frontal systems in attention, working memory, cognitive control, and the integration of internally generated models with ongoing processing^43–45^, domains often reported as atypical across the autism spectrum^63,64^.

While we did not advance a priori predictions regarding hemispheric asymmetry, the confinement of the effect to the left parieto-frontal lines is consistent with the functional profile of left fronto-parietal and fronto-temporal networks, preferentially engaged in language, contextual integration, and aspects of social cognition^65,66^, all domains atypical across the autism continuum. While this interpretation remains speculative, the spatial specificity of the effect and motivates targeted follow-up using task paradigms that explicitly recruit left-hemisphere integrative functions.

An additional strength of the present findings is that the traveling-wave effect was not paralleled by group differences in conventional spectral markers. Oscillatory power, aperiodic parameters, and peak alpha frequency showed no matching pattern of group differences, indicating that the observed reorganization was not accompanied by a generic increase or decrease in alpha activity, a shift in the 1/f background, or a displaced alpha peak. What differed across the autism continuum in our data was not the amount of alpha activity, but its directionally organized pattern across the network. This dissociation suggests that traveling-wave analyses capture aspects of large-scale communication not reducible to power-based measures^33,34,67^. Notably, no reciprocal group effect was observed in the gamma range, despite rhythm-based models of cortical communication typically associating gamma activity with feedforward propagation^12^. This spectral selectivity reinforces the view that the directional reorganization associated with autistic traits is specific to the alpha-band feedback channel, rather than reflecting a system-wide rebalancing of feedback and feedforward spectral markers.

Beyond its mechanistic interest, the present finding has relevance as a candidate trait-sensitive neural marker that scales along the autism continuum rather than being confined to diagnostic categories. Dimensional neural markers are increasingly regarded as a promising tool for psychiatric stratification, particularly in conditions like autism where clinical heterogeneity is high and categorical diagnosis captures only part of the phenotypic variability^68,69^. The directional signature described here is derived from a short resting-state EEG recording, does not require task engagement, and can be estimated with a standard electrode montage, features that make it operationally feasible in both research and clinical settings. Although its clinical utility cannot be assessed on the present data, the combination of dimensional sensitivity, spectral selectivity, and dissociation from conventional spectral descriptors provides a meaningful starting point for follow-up studies examining whether the same directional signature tracks symptom severity in clinical ASD, co-varies with behaviorally relevant measures of prior-based processing, or is responsive to neuromodulatory interventions targeting alpha-band feedback signaling.

Several limitations should be acknowledged. First, our sample was defined on the basis of autistic traits in a non-clinical population rather than a clinical diagnosis. Although dimensional approaches are increasingly recognized as informative in autism research^68,69^, it remains an open question whether the traveling-wave profile observed here generalizes to diagnosed ASD, scales with symptom severity, or remains stable across development. Second, the traveling-wave analysis characterized propagation along predefined antero-posterior electrode lines and thus captured a one-dimensional projection of what is likely to be a more complex two-dimensional cortical wave pattern. Complementary full-scalp approaches, such as phase-gradient methods^70^, may reveal additional spatial structure that the present decomposition cannot resolve. Finally, because all measurements were obtained at rest, the present data cannot establish a mechanistic link between the directional architecture described here and the computational operations it is hypothesized to support.

These limitations point to a clear direction for future work. The most direct test of the present interpretive framework would involve measuring the same traveling-wave metric across both rest and task contexts in the same individuals, while explicitly manipulating prior expectations or sensory precision. If the resting-state architecture described here genuinely reflects a predisposition toward altered prior/evidence balance, then individuals with higher autistic traits should not only show reduced backward alpha propagation at rest, but also exhibit attenuated prior-related modulation of the same directional component during behavior. Extending the approach to clinical cohorts and combining it with neuromodulatory protocols targeting fronto-parietal alpha dynamics would further clarify whether the directional signature observed here reflects a stable trait-level bias or a more plastic feature amenable to intervention.

In summary, the present study shows that variation along the autism continuum is associated with a selective reorganization of resting-state traveling-wave directionality in the left parieto-frontal lines. This reorganization was driven primarily by reduced backward-dominant alpha propagation in individuals with higher autistic traits, accompanied by a weaker reciprocal shift toward forward dominance. The effect was anatomically circumscribed, absent in an occipito-central control line set, and dissociable from group differences in oscillatory power, aperiodic structure, and peak alpha frequency. By identifying a directional rather than amplitude-based signature of autistic traits, these findings highlight traveling waves as a promising framework for characterizing how the intrinsic architecture of large-scale cortical communication varies across the autism continuum.

## Conflict of interest

The authors declare no competing financial interests.

## Acknowledgments

A.A. was funded by the European Union under the European Union’s Horizon 2020 research and innovation program (grant agreement No. 101075930). The copyright holder for this is of the author(s) only and does not necessarily reflect those of the European Union or the European Research Council (ERC). Neither the European Union nor the granting authority can be held responsible for them. V.R. is supported by Next Generation EU (NGEU) and funded by the Ministry of the University and Research (MUR), National Recovery and Research Plan (NRRP) PRIN 2022 (grant n 2022H4ZRSN—CUP J53D23008040006): predictive waves in human perception and individual differences along the autism-schizophrenia continuum (D DN. 104 02.02.2022); (grant n. P2022XAKXL—CUP J53D23017340001): Investigating the plasticity of human predictive coding through neuromodulation (D DN. 1409 14.09.2022); Ministerio de Ciencia, Innovación y Universidades, Spain (PID2019-111335 GA-100); L.T. is supported by Bial Foundation (241/24).

## Author Contributions

Luca Tarasi: Conceptualization, Methodology, Investigation, Visualization, Writing - Original Draft, Writing Review and Editing.

Andrea Alamia: Methodology, Supervision, Writing - Original Draft, Writing - Review and Editing. Vincenzo Romei: Conceptualization, Supervision, Writing - Original Draft, Writing - Review and Editing.

## Data and Code Availability Statement

The data and code that support the findings of this study are available from the corresponding author upon reasonable request.

## Methods

### Participants

Two hundred and one healthy young adults took part in the study. The study was conducted in accordance with the Declaration of Helsinki and was approved by the Bioethics Committee of the University of Bologna. All participants provided written informed consent prior to participation, and all data were analyzed anonymously.

Participants were selected from a larger sample of 323 students at the University of Bologna based on autistic traits measured with the Autism Quotient (AQ^46^). The AQ is a self-report questionnaire designed to quantify autistic traits in the general population and includes five subscales: Social Skill, Communication, Attention Switching, Imagination, and Attention to Detail. Responses were scored using the original dichotomous method (0/1). Following the hierarchical formulation proposed by Hoekstra et al.^47^, we computed the AQ Social Interaction factor by combining the Imagination, Communication, Social Skill, and Attention Switching subscales. This decision was supported by previous studies demonstrating that these subscales share common associations with cognitive and neural features^9^. In contrast, the Attention to Detail dimension deviates significantly from the other scales, showing a strong confound with other personality traits^71–73^. Participants were then divided into two age-matched groups based on the lower and upper terciles of the AQ Social Interaction distribution. The Low-AQ group (N = 95) included participants scoring below the first tertile (score < 10; mean = 6.67 ± 0.20), whereas the High-AQ group (N = 106) included participants scoring above the third tertile (score > 16; mean = 19.82 ± 0.37). The two groups did not differ in age (High AQ: 23.39 ± 0.30 years; Low AQ: 22.87 ± 0.30 years; t (199) = 0.96, p = .338, BF = .237) and Gender (High AQ: 63 Female, Low AQ: 68 Female; X^2^ = 3.26, p = .071, BF = 0.92).

### EEG acquisition and preprocessing

Participants were seated comfortably in a dimly lit room. Electroencephalographic activity (EEG) was recorded at rest for 2 minutes while participants kept their eyes closed. A set of 64 electrodes, mounted according to the international 10–10 system, was used. EEG signals were referenced to the vertex (FCz) and impedances were maintained below 10 kΩ. Data was acquired at a sampling rate of 1000 Hz. EEG data were processed offline using custom MATLAB scripts (version R2020b) and the EEGLAB toolbox. The EEG recordings were first filtered in the 0.5–70 Hz frequency band. Signals were then visually inspected, and noisy channels were identified and spherically interpolated. Following bad channel correction, we re-referenced the EEG recordings to the mean of all electrodes. Lastly, Independent Component Analysis (ICA) was applied to remove EEG artifacts associated with eye blinks and movements.

### Traveling waves analysis

We quantified large-scale traveling waves during resting-state EEG using a 2D-FFT approach adapted from previous work^17,74,75^, with one key modification: rather than restricting the analysis to a few predefined long-range axes, we estimated wave propagation along multiple antero-posterior scalp lines to provide broader sampling of posterior-to-anterior trajectories. Traveling waves were computed along eight predefined parieto-frontal lines forming the primary line set, four per hemisphere, each composed of five electrodes ordered from posterior to anterior. These lines were distributed from lateral to medial positions within each hemisphere (Figure 1). In the left hemisphere, the four parieto-frontal lines were: L1 (TP7-T7-FT7-F7-AF7), L2 (CP5-C5-FC5-F5-AF7), L3 (CP3-C3-FC3-F3-AF3), and L4 (CP1-C1-FC1-F1-AF3). In the right hemisphere, the corresponding lines were: R1 (CP2-C2-FC2-F2-AF4), R2 (CP4-C4-FC4-F4-AF4), R3 (CP6-C6-FC6-F6-AF8), and R4 (TP8-T8-FT8-F8-AF8). As an anatomical control, we defined a second set of eight occipito-central lines, again arranged from posterior to anterior and matched in number and structure to the main set, but sampling a more posterior scalp territory. In the left hemisphere, these were cL1 (O1-PO7-P7-TP7-T7), cL2 (O1-PO3-P5-CP5-C5), cL3 (O1-PO3-P3-CP3-C3), and cL4 (O1-PO3-P1-CP1-C1). In the right hemisphere, the corresponding lines were cR1 (O2-PO4-P2-CP2-C2), cR2 (O2-PO4-P4-CP4-C4), cR3 (O2-PO8-P6-CP6-C6), and cR4 (O2-PO8-P8-TP8-T8). The parieto-frontal set constituted our primary region of interest, while the occipito-central set was analyzed as an anatomical control to verify that any group effect observed in the parieto-frontal set did not reflect global, network-unspecific variability (e.g., diffuse spectral or signal-to-noise differences between AQ groups) that would propagate across the scalp.

Continuous EEG data were segmented into 1-s non-overlapping windows. For each window and electrode line, signals from the five electrodes were arranged into a 2D matrix with time samples on one dimension and electrode position on the other, yielding a time × space map. We then computed the 2D Fast Fourier Transform (2D-FFT) of each map. Directional wave components were quantified from the 2D-FFT power distribution: power in the two relevant quadrants indexed waves propagating in opposite directions along the posterior→anterior electrode ordering, hereafter referred to as forward (FW) and backward (BW) waves. For each frequency bin within 2–45 Hz, we extracted the maximum power within the forward and backward quadrants, producing frequency-resolved spectra for FW and BW waves. To control for non-directional spectral power while removing consistent spatial propagation information, we repeated the same procedure after randomly shuffling the electrode order within each line (preserving the same five signals). This generated surrogate spectra (FW_SS and BW_SS) that preserved power but disrupted directional wave structure. Directional wave strength was expressed in decibels as:

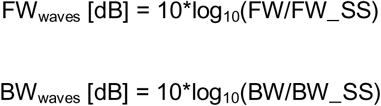

For each electrode line and direction, wave amplitudes were averaged within four canonical frequency bands: theta (4–7 Hz), alpha (8–13 Hz), beta (14–25 Hz), and gamma (26–45 Hz). To summarize directional balance at the single-subject level, we computed a directionality index (DI) as the difference between the band-averaged FW and BW values after surrogate normalization:

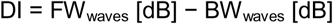

Positive DI values indicate relatively greater FW dominance, negative values relatively greater BW dominance, and values close to zero a more balanced directional profile. As a robust analysis, we also recomputed directional balance using a direct non-surrogate-normalized metric defined as DI = 10 × log10(FW / BW), and repeated the same omnibus and hierarchical follow-up analyses on this alternative measure.

### Statistical analysis of traveling-wave directionality

The primary analysis focused on the parieto-frontal line set. Group differences in DI were tested using a linear mixed-effects model including Band (theta, alpha, beta, gamma), Hemisphere (left, right), and Line Position (1–4, from lateral to medial) as within-subject factors, Group (Low AQ, High AQ) as a between-subject factor, and Subject as a random intercept. The same model structure was then applied to the occipito-central control set. Models were estimated by restricted maximum likelihood with effects coding, and fixed effects were evaluated using F-tests with Satterthwaite degrees of freedom. Significant omnibus interactions were followed up hierarchically. Specifically, when the Group × Band × Hemisphere interaction was significant, we first tested whether this pattern further depended on the Line Position. If the four-way interaction involving Line Position was not significant, DI values were averaged across line positions within subject and the Group × Band interaction was then tested separately within each hemisphere using linear mixed-effects models with Subject as a random intercept. Band-specific between-group contrasts were subsequently performed only within the hemisphere showing a significant Group × Band interaction, and p-values were corrected across bands using the Holm–Bonferroni procedure. To determine whether the critical DI effect was driven primarily by the forward or backward component, we decomposed the relevant Band × Hemisphere effect into FW and BW amplitudes and analyzed them using repeated-measures ANOVA with Direction (FW, BW) as a within-subject factor and Group as a between-subject factor. Significant Group × Direction interactions were followed up with independent samples t-tests on FW and BW separately. For these contrasts, we report both frequentist statistics and Bayes factors (BF10). Pearson correlations between FW and BW were also computed as a descriptive measure of reciprocal coupling, and between-group differences in correlation strength were assessed with Fisher’s r-to-z transformation. Cohen’s d values for key contrasts are reported together with bootstrap 95% confidence intervals based on 1000 resamples. The direct non-surrogate-normalized metric was analyzed with the same omnibus and hierarchical follow-up strategy as the main surrogate-normalized DI.

### Control analyses on spectral features

We next examined whether the group difference in traveling-wave directionality was accompanied by broader group differences in conventional spectral features derived from the same line spectra used in the TW analysis. For each subject and line, single-window spectra were averaged across time to yield one amplitude spectrum per subject and line. All control analyses were computed separately for the parieto-frontal and occipito-central line sets.

### Band-limited power

Amplitude spectra were squared to obtain power, converted to decibels (10 × log10), and averaged within the same four canonical frequency bands used in the TW analysis. Group differences were tested with the same linear mixed-effects structure used for the DI omnibus model, namely Group × Band × Hemisphere × Line Position with Subject as a random intercept.

### Aperiodic component

The aperiodic (1/f) component of each line spectrum was estimated using the FOOOF algorithm^48^ via its Python implementation called from MATLAB. FOOOF was fitted over the 1–40 Hz range using the fixed aperiodic mode (offset + exponent), peak width limits of 1–12 Hz, a maximum of 6 peaks, a minimum peak height of 0.1, and a peak threshold of 2. Fit quality was quantified using the coefficient of determination (R²) and model error. For each subject and line, we extracted the aperiodic exponent and offset. Group differences in these parameters were tested with linear mixed-effects models including Group, Hemisphere, and Line Position as fixed factors and Subject as a random intercept.

### Peak alpha frequency

Peak alpha frequency (PAF) was derived from the FOOOF periodic component and defined as the frequency of the highest-power periodic peak falling within 7–13 Hz after removal of the aperiodic background. When multiple alpha peaks were detected, the one with the highest peak power was retained. When no peak was detected in the alpha range, PAF was coded as missing. Group differences in PAF were tested with the same Group × Hemisphere × Line Position mixed-effects structure used for the aperiodic parameters. Because PAF is only defined when an alpha peak is detected, analyses were performed on valid line-wise observations only.

## Notes

### Competing Interest Statement

The authors have declared no competing interest.

## Reference

1. Ippolito, G. et al. The Role of Alpha Oscillations among the Main Neuropsychiatric Disorders in the Adult and Developing Human Brain: Evidence from the Last 10 Years of Research. Biomedicines 10, 3189 (2022).

2. Kessler, K., Seymour, R. A. & Rippon, G. Brain oscillations and connectivity in autism spectrum disorders (ASD): new approaches to methodology, measurement and modelling. Neuroscience & Biobehavioral Reviews 71, 601–620 (2016).

3. Di Martino, A. et al. The autism brain imaging data exchange: towards a large-scale evaluation of the intrinsic brain architecture in autism. Mol Psychiatry 19, 659–667 (2014).

4. Holiga, Š. et al. Patients with autism spectrum disorders display reproducible functional connectivity alterations. Sci Transl Med 11, eaat9223 (2019).

5. Just, M. A., Keller, T. A., Malave, V. L., Kana, R. K. & Varma, S. Autism as a neural systems disorder: a theory of frontal-posterior underconnectivity. Neurosci Biobehav Rev 36, 1292–1313 (2012).

6. Clark, A. Whatever next? Predictive brains, situated agents, and the future of cognitive science. Behav Brain Sci 36, 181–204 (2013).

7. Friston, K. & Kiebel, S. Predictive coding under the free-energy principle. Philos Trans R Soc Lond B Biol Sci 364, 1211–1221 (2009).

8. Andersen, B. P. Autistic-Like Traits and Positive Schizotypy as Diametric Specializations of the Predictive Mind. Perspect Psychol Sci 17, 1653–1672 (2022).

9. Tarasi, L. et al. Predictive waves in the autism-schizophrenia continuum: A novel biobehavioral model. Neurosci Biobehav Rev 132, 1–22 (2022).

10. Pellicano, E. & Burr, D. When the world becomes ‘too real’: a Bayesian explanation of autistic perception. Trends Cogn Sci 16, 504–510 (2012).

11. Lawson, R. P., Rees, G. & Friston, K. J. An aberrant precision account of autism. Front Hum Neurosci 8, 302 (2014).

12. Bastos, A. M. et al. Canonical microcircuits for predictive coding. Neuron 76, 695–711 (2012).

13. Bastos, A. M., Lundqvist, M., Waite, A. S., Kopell, N. & Miller, E. K. Layer and rhythm specificity for predictive routing. Proceedings of the National Academy of Sciences 117, 31459–31469 (2020).

14. Chao, Z. C., Takaura, K., Wang, L., Fujii, N. & Dehaene, S. Large-Scale Cortical Networks for Hierarchical Prediction and Prediction Error in the Primate Brain. Neuron 100, 1252–1266.e3 (2018).

15. Michalareas, G. et al. Alpha-Beta and Gamma Rhythms Subserve Feedback and Feedforward Influences among Human Visual Cortical Areas. Neuron 89, 384–397 (2016).

16. Tarasi, L., di Pellegrino, G. & Romei, V. Are you an empiricist or a believer? Neural signatures of predictive strategies in humans. Progress in Neurobiology 219, 102367 (2022).

17. Alamia, A., Terral, L., D’ambra, M. R. & VanRullen, R. Distinct roles of forward and backward alpha-band waves in spatial visual attention. eLife 12, e85035 (2023).

18. Bollimunta, A., Mo, J., Schroeder, C. E. & Ding, M. Neuronal Mechanisms and Attentional Modulation of Corticothalamic Alpha Oscillations. J Neurosci 31, 4935–4943 (2011).

19. Bonnefond, M. & Jensen, O. The role of alpha oscillations in resisting distraction. Trends Cogn Sci 29, 368–379 (2025).

20. Frisoni, M., Tarasi, L., Borgomaneri, S. & Romei, V. The relationship between individual alpha frequency and time perception: Testing the internal clock versus the sampling rate hypothesis. Cortex 192, 183–195 (2025).

21. Palva, S. & Palva, J. M. New vistas for alpha-frequency band oscillations. Trends Neurosci 30, 150– 158 (2007).

22. Pascucci, D. et al. EEG brain waves and alpha rhythms: Past, current and future direction. Neuroscience & Biobehavioral Reviews 176, 106288 (2025).

23. Romei, V. & Tarasi, L. Alpha frequency shapes perceptual sensitivity by modulating optimal phase likelihood. Nat Commun 17, 3384 (2026).

24. Santoni, A. et al. Bifocal alpha-band tACS modulates temporal sampling in visual perception. NeuroImage 320, 121474 (2025).

25. Tarasi, L. & Romei, V. Individual Alpha Frequency Contributes to the Precision of Human Visual Processing. J Cogn Neurosci 36, 602–613 (2024).

26. Jensen, O., Bonnefond, M., Marshall, T. R. & Tiesinga, P. Oscillatory mechanisms of feedforward and feedback visual processing. Trends in Neurosciences 38, 192–194 (2015).

27. van Kerkoerle, T. et al. Alpha and gamma oscillations characterize feedback and feedforward processing in monkey visual cortex. Proc Natl Acad Sci U S A 111, 14332–14341 (2014).

28. Coll, M.-P., Whelan, E., Catmur, C. & Bird, G. Autistic traits are associated with atypical precision-weighted integration of top-down and bottom-up neural signals. Cognition 199, 104236 (2020).

29. Cook, J. L., Barbalat, G. & Blakemore, S.-J. Top-down modulation of the perception of other people in schizophrenia and autism. Front. Hum. Neurosci. 6, (2012).

30. Knight, E. J. et al. Severely Attenuated Visual Feedback Processing in Children on the Autism Spectrum. J Neurosci 43, 2424–2438 (2023).

31. Ursino, M. et al. Bottom-up vs. top-down connectivity imbalance in individuals with high-autistic traits: An electroencephalographic study. Front. Syst. Neurosci. 16, (2022).

32. Tarasi, L., Magosso, E., Ricci, G., Ursino, M. & Romei, V. The Directionality of Fronto-Posterior Brain Connectivity Is Associated with the Degree of Individual Autistic Traits. Brain Sciences 11, 1443 (2021).

33. Alamia, A. & VanRullen, R. Alpha oscillations and traveling waves: Signatures of predictive coding? PLOS Biology 17, e3000487 (2019).

34. Muller, L., Chavane, F., Reynolds, J. & Sejnowski, T. J. Cortical travelling waves: mechanisms and computational principles. Nat Rev Neurosci 19, 255–268 (2018).

35. Petras, K., Grabot, L. & Dugué, L. Locally Induced Traveling Waves Generate Globally Observable Traveling Waves. J Neurosci 45, e0089252025 (2025).

36. Sato, T. K., Nauhaus, I. & Carandini, M. Traveling waves in visual cortex. Neuron 75, 218–229 (2012).

37. Zhang, H., Watrous, A. J., Patel, A. & Jacobs, J. Theta and Alpha Oscillations Are Traveling Waves in the Human Neocortex. Neuron 98, 1269–1281.e4 (2018).

38. Fox, M. D. & Raichle, M. E. Spontaneous fluctuations in brain activity observed with functional magnetic resonance imaging. Nat Rev Neurosci 8, 700–711 (2007).

39. Raichle, M. E. The brain’s default mode network. Annu Rev Neurosci 38, 433–447 (2015).

40. Tarasi, L., Romanazzi, D., Pasini, A. & Romei, V. Delusion-like thinking is associated with lower individual alpha peak frequency. Schizophr 11, 76 (2025).

41. Trajkovic, J. et al. Aberrant Functional Connectivity and Brain Network Organization in High-Schizotypy Individuals: An Electroencephalography Study. Schizophr Bull 51, 1266–1281 (2025).

42. Preacher, K. J., Rucker, D. D., MacCallum, R. C. & Nicewander, W. A. Use of the extreme groups approach: a critical reexamination and new recommendations. Psychol Methods 10, 178–192 (2005).

43. Corbetta, M. & Shulman, G. L. Control of goal-directed and stimulus-driven attention in the brain. Nat Rev Neurosci 3, 201–215 (2002).

44. Duncan, J. The multiple-demand (MD) system of the primate brain: mental programs for intelligent behaviour. Trends in Cognitive Sciences 14, 172–179 (2010).

45. Summerfield, C. & de Lange, F. P. Expectation in perceptual decision making: neural and computational mechanisms. Nat Rev Neurosci 15, 745–756 (2014).

46. Baron-Cohen, S., Wheelwright, S., Skinner, R., Martin, J. & Clubley, E. The autism-spectrum quotient (AQ): evidence from Asperger syndrome/high-functioning autism, males and females, scientists and mathematicians. J Autism Dev Disord 31, 5–17 (2001).

47. Hoekstra, R. A., Bartels, M., Cath, D. C. & Boomsma, D. I. Factor structure, reliability and criterion validity of the Autism-Spectrum Quotient (AQ): a study in Dutch population and patient groups. J Autism Dev Disord 38, 1555–1566 (2008).

48. Donoghue, T. et al. Parameterizing neural power spectra into periodic and aperiodic components. Nat Neurosci 23, 1655–1665 (2020).

49. Murias, M., Webb, S. J., Greenson, J. & Dawson, G. Resting state cortical connectivity reflected in EEG coherence in individuals with autism. Biol Psychiatry 62, 270–273 (2007).

50. Keehn, B., Westerfield, M., Müller, R.-A. & Townsend, J. Autism, Attention, and Alpha Oscillations: An Electrophysiological Study of Attentional Capture. Biol Psychiatry Cogn Neurosci Neuroimaging 2, 528–536 (2017).

51. Yuk, V., Urbain, C., Anagnostou, E. & Taylor, M. J. Frontoparietal Network Connectivity During an N-Back Task in Adults With Autism Spectrum Disorder. Front. Psychiatry 11, (2020).

52. Seymour, R. A., Rippon, G., Gooding-Williams, G., Schoffelen, J. M. & Kessler, K. Dysregulated oscillatory connectivity in the visual system in autism spectrum disorder. Brain 142, 3294–3305 (2019).

53. Alamia, A., Schwenk, J. C. B., Wagemans, J. & Sapey-Triomphe, L.-A. Oscillatory traveling waves during visual entrainment in autistic and neurotypical adults. 2026.01.29.702587 Preprint at 10.64898/2026.01.29.702587 (2026).

54. Tarasi, L. et al. Oscillatory signatures of monitoring and anticipatory strategies for probabilistic vs deterministic cues. Imaging Neuroscience 3, imag_a_00496 (2025).

55. Arnal, L. H. & Giraud, A.-L. Cortical oscillations and sensory predictions. Trends Cogn Sci 16, 390– 398 (2012).

56. Halgren, M. et al. The generation and propagation of the human alpha rhythm. Proceedings of the National Academy of Sciences 116, 23772–23782 (2019).

57. Karvelis, P., Seitz, A. R., Lawrie, S. M. & Seriès, P. Autistic traits, but not schizotypy, predict increased weighting of sensory information in Bayesian visual integration. Elife 7, e34115 (2018).

58. Palmer, C. J., Lawson, R. P. & Hohwy, J. Bayesian approaches to autism: Towards volatility, action, and behavior. Psychological Bulletin 143, 521–542 (2017).

59. Tarasi, L., et al. Preparing to act follows Bayesian inference rules. iScience 28, (2025).

60. Van de Cruys, S. et al. Precise minds in uncertain worlds: predictive coding in autism. Psychol Rev 121, 649–675 (2014).

61. Tarasi, L., Martelli, M. E., Bortoletto, M., di Pellegrino, G. & Romei, V. Neural Signatures of Predictive Strategies Track Individuals Along the Autism-Schizophrenia Continuum. Schizophr Bull 49, 1294–1304 (2023).

62. Tarasi, L., Alamia, A. & Romei, V. Backward alpha band oscillations shape perceptual bias under probabilistic cues. Commun Biol https://doi.org/10.1038/s42003-026-09559-1 (2026) doi:10.1038/s42003-026-09559-1.

63. Hill, E. L. Executive dysfunction in autism. Trends Cogn Sci 8, 26–32 (2004).

64. Solomon, M. et al. The neural substrates of cognitive control deficits in autism spectrum disorders. Neuropsychologia 47, 2515–2526 (2009).

65. Eyler, L. T., Pierce, K. & Courchesne, E. A failure of left temporal cortex to specialize for language is an early emerging and fundamental property of autism. Brain 135, 949–960 (2012).

66. Lombardo, M. V. et al. Different functional neural substrates for good and poor language outcome in autism. Neuron 86, 567–577 (2015).

67. Davis, Z. W., Muller, L., Martinez-Trujillo, J., Sejnowski, T. & Reynolds, J. H. Spontaneous travelling cortical waves gate perception in behaving primates. Nature 587, 432–436 (2020).

68. Constantino, J. N. & Todd, R. D. Autistic traits in the general population: a twin study. Arch Gen Psychiatry 60, 524–530 (2003).

69. Ruzich, E. et al. Measuring autistic traits in the general population: a systematic review of the Autism-Spectrum Quotient (AQ) in a nonclinical population sample of 6,900 typical adult males and females. Mol Autism 6, 2 (2015).

70. Das, A. et al. Spontaneous neuronal oscillations in the human insula are hierarchically organized traveling waves. eLife 11, e76702 (2022).

71. Dinsdale, N. L., Hurd, P. L., Wakabayashi, A., Elliot, M. & Crespi, B. J. How Are Autism and Schizotypy Related? Evidence from a Non-Clinical Population. PLOS ONE 8, e63316 (2013).

72. Nenadić, I. et al. Subclinical schizotypal vs. autistic traits show overlapping and diametrically opposed facets in a non-clinical population. Schizophrenia Research 231, 32–41 (2021).

73. Tarasi, L., Borgomaneri, S. & Romei, V. Antivax attitude in the general population along the autism-schizophrenia continuum and the impact of socio-demographic factors. Front. Psychol. 14, (2023).

74. Alamia, A. et al. Oscillatory traveling waves provide evidence for predictive coding abnormalities in schizophrenia. Biological Psychiatry https://doi.org/10.1016/j.biopsych.2024.11.014 (2024) doi:10.1016/j.biopsych.2024.11.014.

75. Tarasi, L., Alamia, A. & Romei, V. Perceptual Bias in Motion Discrimination is Related to Asymmetric Interhemispheric Alpha Traveling Waves. Advanced Science 12, e14623 (2025).

