## Supplementary materials for "Reduced Backward Alpha Propagation at Rest Marks the Autism Continuum"

**Table S1. Omnibus linear mixed-effects model on parieto-frontal directionality index (DI = FW − BW).**

| **Term** | **F** | **df1** | **df2** | **p** |
| --- | --- | --- | --- | --- |
| Group | 3.409 | 1 | 199 | .066 |
| **Band** | **113.290** | **3** | **6169** | **< .001** |
| **Hemisphere** | **14.861** | **1** | **6169** | **< .001** |
| **LinePos** | **51.983** | **3** | **6169** | **< .001** |
| **Group × Band** | **7.589** | **3** | **6169** | **< .001** |
| Group × Hemisphere | 0.547 | 1 | 6169 | .460 |
| Band × Hemisphere | 2.146 | 3 | 6169 | .092 |
| Group × LinePos | 0.458 | 3 | 6169 | .712 |
| **Band × LinePos** | **28.923** | **9** | **6169** | **< .001** |
| Hemisphere × LinePos | 0.630 | 3 | 6169 | .596 |
| **Group × Band × Hemisphere** | **3.642** | **3** | **6169** | **.012** |
| Group × Band × LinePos | 1.426 | 9 | 6169 | .171 |
| Group × Hemisphere × LinePos | 0.147 | 3 | 6169 | .932 |
| Band × Hemisphere × LinePos | 0.604 | 9 | 6169 | .795 |
| Group × Band × Hemisphere × LinePos | 0.619 | 9 | 6169 | .782 |

*Note.* Fixed-effects ANOVA marginal tests with Satterthwaite degrees of freedom from the LME NetDir ~ Group × Band × Hemisphere × LinePos + (1|Subject), fitted via REML with effects coding. The intercept term is omitted for readability.

**Table S2. Hierarchical follow-up of the parieto-frontal DI interaction: Group × Band tested separately within each hemisphere.**

| **Hemisphere** | **Term** | **F** | **df1** | **df2** | **p** |
| --- | --- | --- | --- | --- | --- |
| Left | Group | 3.673 | 1 | 199 | .057 |
| **Left** | **Band** | **22.165** | **3** | **597** | **< .001** |
| **Left** | **Group × Band** | **4.150** | **3** | **597** | **.006** |
| Right | Group | 1.925 | 1 | 199 | .167 |
| **Right** | **Band** | **28.870** | **3** | **597** | **< .001** |
| Right | Group × Band | 0.691 | 3 | 597 | .558 |

*Note.* Follow-up LMEs performed after averaging DI across line positions within subject, because the omnibus Group × Band × Hemisphere × LinePos interaction was null. Each model tested NetDir ~ Group × Band + (1|Subject) separately within the left and right hemispheres.

**Table S3. Band-specific group contrasts on parieto-frontal DI within the left hemisphere.**

| **Band** | **t** | **df** | **p** | **p (Holm)** | **Cohen's d** | **BF₁₀** |
| --- | --- | --- | --- | --- | --- | --- |
| Theta | -0.32 | 199 | .750 | 1.000 | -0.05 | 0.16 |
| **Alpha** | **-2.73** | **199** | **.007** | **.028** | **-0.38** | **4.77** |
| Beta | -0.49 | 199 | .622 | 1.000 | -0.07 | 0.17 |
| Gamma | 0.09 | 199 | .928 | 1.000 | 0.01 | 0.15 |

*Note.* Independent-samples t-tests (df = 199) comparing low-AQ and high-AQ groups on DI averaged across line positions within each band of the left hemisphere. Holm correction was applied across the four band-level contrasts. Cohen’s d follows the convention LOW − HIGH; negative values indicate higher DI in the high-AQ group. BF₁₀ quantifies evidence for a group difference relative to the null.

**Table S4. Omnibus linear mixed-effects model on DI = [10 * log10 (FW/BW)].**

| **Term** | **F** | **df1** | **df2** | **p** |
| --- | --- | --- | --- | --- |
| Group | 3.698 | 1 | 199 | .056 |
| **Band** | **116.380** | **3** | **6169** | **< .001** |
| **Hemisphere** | **13.375** | **1** | **6169** | **< .001** |
| **LinePos** | **50.294** | **3** | **6169** | **< .001** |
| **Group × Band** | **6.836** | **3** | **6169** | **< .001** |
| Group × Hemisphere | 0.676 | 1 | 6169 | .411 |
| Band × Hemisphere | 2.124 | 3 | 6169 | .095 |
| Group × LinePos | 0.236 | 3 | 6169 | .871 |
| **Band × LinePos** | **28.518** | **9** | **6169** | **< .001** |
| Hemisphere × LinePos | 0.894 | 3 | 6169 | .443 |
| **Group × Band × Hemisphere** | **3.006** | **3** | **6169** | **.029** |
| Group × Band × LinePos | 1.506 | 9 | 6169 | .139 |
| Group × Hemisphere × LinePos | 0.254 | 3 | 6169 | .859 |
| Band × Hemisphere × LinePos | 0.575 | 9 | 6169 | .818 |
| Group × Band × Hemisphere × LinePos | 0.618 | 9 | 6169 | .783 |

*Note.* Robustness analysis using a direct directional metric without surrogate normalization: DI = 10·log10(FW/BW). Fixed-effects ANOVA marginal tests with Satterthwaite degrees of freedom from the LME DI ~ Group × Band × Hemisphere × LinePos + (1|Subject), fitted via REML with effects coding.

**Table S5. Hierarchical follow-up of the direct-DI interaction: Group × Band tested separately within each hemisphere.**

| **Hemisphere** | **Term** | **F** | **df1** | **df2** | **p** |
| --- | --- | --- | --- | --- | --- |
| **Left** | **Group** | **4.061** | **1** | **199** | **.045** |
| **Left** | **Band** | **22.796** | **3** | **597** | **< .001** |
| **Left** | **Group × Band** | **3.637** | **3** | **597** | **.013** |
| Right | Group | 2.001 | 1 | 199 | .159 |
| **Right** | **Band** | **30.238** | **3** | **597** | **< .001** |
| Right | Group × Band | 0.629 | 3 | 597 | .597 |

*Note.* Follow-up LMEs performed after averaging direct DI across line positions within subject. Each model tested DI ~ Group × Band + (1|Subject) separately within the left and right hemispheres.

**Table S6. Band-specific group contrasts on direct DI within the left hemisphere.**

| **Band** | **t** | **df** | **p** | **p (Holm)** | **Cohen's d** | **BF₁₀** |
| --- | --- | --- | --- | --- | --- | --- |
| Theta | -0.68 | 199 | .499 | 1.000 | -0.10 | 0.19 |
| **Alpha** | **-2.64** | **199** | **.009** | **.036** | **-0.37** | **3.88** |
| Beta | -0.53 | 199 | .600 | 1.000 | -0.07 | 0.17 |
| Gamma | 0.07 | 199 | .945 | 1.000 | 0.01 | 0.15 |

*Note.* Independent-samples t-tests (df = 199) comparing low-AQ and high-AQ groups on direct DI averaged across line positions within each band of the left hemisphere. Holm correction was applied across the four band-level contrasts. Cohen’s d follows the convention LOW − HIGH. BF₁₀ quantifies evidence for a group difference relative to the null.

**Table S7. Omnibus linear mixed-effects model on parieto-frontal band-limited power.**

| **Term** | **F** | **df1** | **df2** | **p** |
| --- | --- | --- | --- | --- |
| Group | 3.406 | 1 | 199 | .066 |
| **Band** | **15179.000** | **3** | **6169** | **< .001** |
| Hemisphere | 3.697 | 1 | 6169 | .055 |
| **LinePos** | **495.050** | **3** | **6169** | **< .001** |
| **Group × Band** | **3.225** | **3** | **6169** | **.022** |
| Group × Hemisphere | 0.640 | 1 | 6169 | .424 |
| Band × Hemisphere | 0.301 | 3 | 6169 | .825 |
| Group × LinePos | 1.303 | 3 | 6169 | .272 |
| **Band × LinePos** | **71.440** | **9** | **6169** | **< .001** |
| Hemisphere × LinePos | 0.608 | 3 | 6169 | .610 |
| Group × Band × Hemisphere | 0.074 | 3 | 6169 | .974 |
| Group × Band × LinePos | 0.309 | 9 | 6169 | .972 |
| Group × Hemisphere × LinePos | 0.316 | 3 | 6169 | .814 |
| Band × Hemisphere × LinePos | 0.141 | 9 | 6169 | .999 |
| Group × Band × Hemisphere × LinePos | 0.033 | 9 | 6169 | 1.000 |

*Note.* Fixed-effects ANOVA marginal tests on the same factorial structure used for the DI omnibus, applied to parieto-frontal band-limited power as a control analysis.

**Table S8. Group contrasts on parieto-frontal band-limited power, per frequency band.**

| **Band** | **Mean LOW** | **Mean HIGH** | **t** | **df** | **p** | **Cohen's d** | **BF₁₀** | **p (Holm)** |
| --- | --- | --- | --- | --- | --- | --- | --- | --- |
| Theta | 56.104 | 55.607 | 1.50 | 199 | .135 | 0.21 | 0.44 | .334 |
| Alpha | 57.196 | 56.373 | 1.60 | 199 | .111 | 0.23 | 0.51 | .334 |
| Beta | 49.582 | 49.013 | 1.75 | 199 | .081 | 0.25 | 0.64 | .325 |
| Gamma | 45.295 | 44.830 | 1.40 | 199 | .164 | 0.20 | 0.38 | .334 |

*Note.* Independent-samples t-tests (df = 199) comparing low-AQ and high-AQ groups on parieto-frontal power averaged across hemispheres and line positions within each band. Holm correction was applied across the four band-level contrasts. No band-level comparison survived correction.

**Table S9. Linear mixed-effects models on parieto-frontal FOOOF-derived spectral parameters.**

| **DV** | **Term** | **F** | **df1** | **df2** | **p** |
| --- | --- | --- | --- | --- | --- |
| Exponent | Group | 0.000 | 1 | 199 | .983 |
| Exponent | Hemisphere | 2.800 | 1 | 1393 | .094 |
| **Exponent** | **LinePos** | **249.990** | **3** | **1393** | **< .001** |
| Exponent | Group × Hemisphere | 0.037 | 1 | 1393 | .847 |
| **Exponent** | **Group × LinePos** | **2.678** | **3** | **1393** | **.046** |
| Exponent | Hemisphere × LinePos | 0.072 | 3 | 1393 | .975 |
| Exponent | Group × Hemisphere × LinePos | 0.144 | 3 | 1393 | .933 |
| Offset | Group | 2.027 | 1 | 199 | .156 |
| Offset | Hemisphere | 0.400 | 1 | 1393 | .527 |
| **Offset** | **LinePos** | **185.860** | **3** | **1393** | **< .001** |
| Offset | Group × Hemisphere | 0.357 | 1 | 1393 | .550 |
| Offset | Group × LinePos | 1.077 | 3 | 1393 | .358 |
| Offset | Hemisphere × LinePos | 0.480 | 3 | 1393 | .697 |
| Offset | Group × Hemisphere × LinePos | 0.521 | 3 | 1393 | .668 |
| PAF | Group | 0.203 | 1 | 194.95 | .653 |
| PAF | Hemisphere | 0.000 | 1 | 1329.4 | .993 |
| PAF | LinePos | 2.210 | 3 | 1329.6 | .085 |
| PAF | Group × Hemisphere | 0.574 | 1 | 1329.4 | .449 |
| PAF | Group × LinePos | 0.738 | 3 | 1329.6 | .529 |
| PAF | Hemisphere × LinePos | 1.387 | 3 | 1329.4 | .245 |
| PAF | Group × Hemisphere × LinePos | 0.064 | 3 | 1329.4 | .979 |

*Note.* Fixed-effects ANOVA marginal tests with Satterthwaite degrees of freedom from LMEs of the form DV ~ Group × Hemisphere × LinePos + (1|Subject) on line-wise FOOOF-derived parameters from the parieto-frontal spectra. PAF denotes the centre frequency of the highest-power periodic component within 7–13 Hz after removal of the aperiodic background. FOOOF fit quality was high (mean R² = 0.981, SD = 0.037; mean fit error = 0.046, SD = 0.021); 68/1608 line-wise PAF estimates were missing (LOW: 32/760, 4.2%; HIGH: 36/848, 4.2%).

**Table S10. Follow-up group contrasts on the parieto-frontal aperiodic exponent by line position.**

| **LinePos** | **Mean LOW** | **Mean HIGH** | **t** | **df** | **p** | **Cohen's d** | **BF₁₀** | **p (Holm)** |
| --- | --- | --- | --- | --- | --- | --- | --- | --- |
| 1 | 1.085 | 1.064 | 0.67 | 199 | .505 | 0.09 | 0.19 | 1.000 |
| 2 | 1.075 | 1.067 | 0.28 | 199 | .779 | 0.04 | 0.16 | 1.000 |
| 3 | 1.135 | 1.150 | -0.61 | 199 | .539 | -0.09 | 0.18 | 1.000 |
| 4 | 1.244 | 1.256 | -0.57 | 199 | .570 | -0.08 | 0.18 | 1.000 |

*Note.* Follow-up independent-samples t-tests (df = 199) comparing low-AQ and high-AQ groups on the parieto-frontal aperiodic exponent after averaging across hemispheres within each line position. Holm correction was applied across the four line-position contrasts. No line-specific contrast survived correction.

**Table S11. Omnibus linear mixed-effects model on occipito-central directionality index.**

| **Term** | **F** | **df1** | **df2** | **p** |
| --- | --- | --- | --- | --- |
| Group | 0.017 | 1 | 199 | .898 |
| **Band** | **336.190** | **3** | **6169** | **< .001** |
| **Hemisphere** | **44.613** | **1** | **6169** | **< .001** |
| **LinePos** | **37.143** | **3** | **6169** | **< .001** |
| Group × Band | 1.396 | 3 | 6169 | .242 |
| Group × Hemisphere | 0.009 | 1 | 6169 | .925 |
| **Band × Hemisphere** | **15.303** | **3** | **6169** | **< .001** |
| Group × LinePos | 1.915 | 3 | 6169 | .125 |
| **Band × LinePos** | **21.001** | **9** | **6169** | **< .001** |
| **Hemisphere × LinePos** | **6.681** | **3** | **6169** | **< .001** |
| Group × Band × Hemisphere | 0.129 | 3 | 6169 | .943 |
| Group × Band × LinePos | 0.186 | 9 | 6169 | .996 |
| Group × Hemisphere × LinePos | 2.127 | 3 | 6169 | .095 |
| Band × Hemisphere × LinePos | 1.362 | 9 | 6169 | .199 |
| Group × Band × Hemisphere × LinePos | 0.576 | 9 | 6169 | .818 |

*Note.* Same omnibus model structure as Table S1, fitted to the occipito-central control line set. No group-related term approached significance.

**Table S12. Omnibus linear mixed-effects model on occipito-central band-limited power.**

| **Term** | **F** | **df1** | **df2** | **p** |
| --- | --- | --- | --- | --- |
| Group | 2.788 | 1 | 199 | .097 |
| **Band** | **19939.000** | **3** | **6169** | **< .001** |
| **Hemisphere** | **40.721** | **1** | **6169** | **< .001** |
| **LinePos** | **274.010** | **3** | **6169** | **< .001** |
| **Group × Band** | **3.522** | **3** | **6169** | **.014** |
| Group × Hemisphere | 0.092 | 1 | 6169 | .762 |
| Band × Hemisphere | 1.938 | 3 | 6169 | .121 |
| Group × LinePos | 0.604 | 3 | 6169 | .612 |
| **Band × LinePos** | **22.711** | **9** | **6169** | **< .001** |
| **Hemisphere × LinePos** | **2.680** | **3** | **6169** | **.045** |
| Group × Band × Hemisphere | 0.143 | 3 | 6169 | .934 |
| Group × Band × LinePos | 0.064 | 9 | 6169 | 1.000 |
| Group × Hemisphere × LinePos | 0.611 | 3 | 6169 | .608 |
| Band × Hemisphere × LinePos | 0.375 | 9 | 6169 | .947 |
| Group × Band × Hemisphere × LinePos | 0.034 | 9 | 6169 | 1.000 |

*Note.* Fixed-effects ANOVA marginal tests on the same factorial structure used for the DI omnibus, applied to occipito-central band-limited power as a control analysis.

**Table S13. Group contrasts on occipito-central band-limited power, per frequency band.**

| **Band** | **Mean LOW** | **Mean HIGH** | **t** | **df** | **p** | **Cohen's d** | **BF₁₀** | **p (Holm)** |
| --- | --- | --- | --- | --- | --- | --- | --- | --- |
| Theta | 56.581 | 55.981 | 1.53 | 199 | .126 | 0.22 | 0.46 | .379 |
| Alpha | 59.665 | 58.806 | 1.41 | 199 | .161 | 0.20 | 0.39 | .379 |
| Beta | 50.411 | 49.785 | 1.69 | 199 | .092 | 0.24 | 0.59 | .368 |
| Gamma | 44.884 | 44.444 | 1.27 | 199 | .205 | 0.18 | 0.33 | .379 |

*Note.* Independent-samples t-tests (df = 199) comparing low-AQ and high-AQ groups on occipito-central power averaged across hemispheres and line positions within each band. Holm correction was applied across the four band-level contrasts. No band-level comparison survived correction.

**Table S14. Linear mixed-effects models on occipito-central FOOOF-derived spectral parameters.**

| **DV** | **Term** | **F** | **df1** | **df2** | **p** |
| --- | --- | --- | --- | --- | --- |
| Exponent | Group | 0.021 | 1 | 199 | .884 |
| Exponent | Hemisphere | 1.958 | 1 | 1393 | .162 |
| **Exponent** | **LinePos** | **231.690** | **3** | **1393** | **< .001** |
| Exponent | Group × Hemisphere | 1.914 | 1 | 1393 | .167 |
| Exponent | Group × LinePos | 0.264 | 3 | 1393 | .851 |
| Exponent | Hemisphere × LinePos | 1.945 | 3 | 1393 | .121 |
| Exponent | Group × Hemisphere × LinePos | 0.104 | 3 | 1393 | .958 |
| Offset | Group | 2.135 | 1 | 199 | .146 |
| **Offset** | **Hemisphere** | **48.271** | **1** | **1393** | **< .001** |
| **Offset** | **LinePos** | **87.410** | **3** | **1393** | **< .001** |
| Offset | Group × Hemisphere | 0.816 | 1 | 1393 | .367 |
| Offset | Group × LinePos | 0.318 | 3 | 1393 | .812 |
| Offset | Hemisphere × LinePos | 0.829 | 3 | 1393 | .478 |
| Offset | Group × Hemisphere × LinePos | 1.242 | 3 | 1393 | .293 |
| PAF | Group | 0.191 | 1 | 197.41 | .663 |
| PAF | Hemisphere | 2.096 | 1 | 1368.6 | .148 |
| **PAF** | **LinePos** | **9.523** | **3** | **1368.5** | **< .001** |
| PAF | Group × Hemisphere | 0.652 | 1 | 1368.6 | .420 |
| PAF | Group × LinePos | 0.371 | 3 | 1368.5 | .774 |
| PAF | Hemisphere × LinePos | 0.444 | 3 | 1368.5 | .722 |
| PAF | Group × Hemisphere × LinePos | 0.940 | 3 | 1368.5 | .421 |

*Note.* Fixed-effects ANOVA marginal tests with Satterthwaite degrees of freedom from LMEs of the form DV ~ Group × Hemisphere × LinePos + (1|Subject) on line-wise FOOOF-derived parameters from the occipito-central control spectra. FOOOF fit quality was high (mean R² = 0.986, SD = 0.014; mean fit error = 0.051, SD = 0.021); 25/1608 line-wise PAF estimates were missing (LOW: 11/760, 1.4%; HIGH: 14/848, 1.7%).

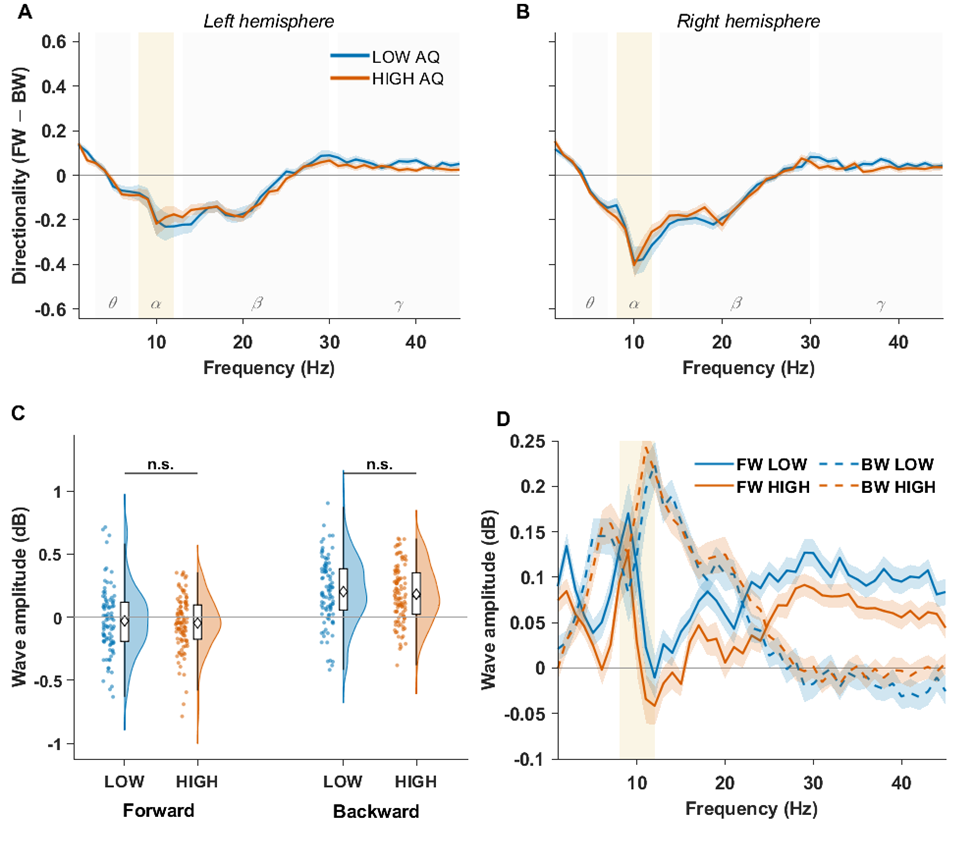

***Figure S1. Occipito-Central traveling-wave directionality in low vs high AQ groups****.*

*(A, B) Directionality index (DI; computed as the difference between FW and BW wave amplitudes after surrogate normalization, in dB) as a function of frequency, averaged across the four occipito-central lines of the left (A) and right (B) hemisphere. Solid lines show group means for low-AQ (blue) and high-AQ (orange) participants; shaded areas denote ± 1 SEM. Canonical frequency bands (θ, α, β, γ) are indicated by background shading, with the alpha band highlighted. Positive DI values reflect forward-dominant propagation, negative values backward-dominant propagation. The omnibus LME on occipito-central DI showed no effect of Group (F(1, 199) = 0.02, p = .898) and, crucially, no Group × Band (F(3, 6169) = 1.40, p = .242), Group × Hemisphere (F(1, 6169) = 0.01, p = .925), or Group × Band × Hemisphere interaction (F(3, 6169) = 0.13, p = .943). No interaction involving Group approached significance, including the Group × Band × Hemisphere × LinePos term (F(9, 6169) = 0.58, p = .818) (full statistics in Supplementary Table S11). (C) Illustrative decomposition of left-hemisphere alpha FW and BW components in the occipito-central control line set, shown to mirror the layout of Figure 2. Raincloud plots show the distribution of single-subject FW and BW amplitudes for low-AQ (blue) and high-AQ (orange) participants, combining a half-violin kernel density estimate, jittered individual data points, a compact boxplot (median, interquartile range, whiskers at 1.5 × IQR), and the group mean (white diamond). No statistical FW/BW decomposition was performed because the omnibus model revealed no reliable group-related effect. (D) Frequency-resolved forward and backward wave spectra at a representative left-hemisphere occipito-central line, shown for visualization purposes. Solid lines indicate forward waves, dashed lines backward waves; colors code AQ group as in panels A–C. Shaded areas denote ± 1 SEM. The alpha band is highlighted.*

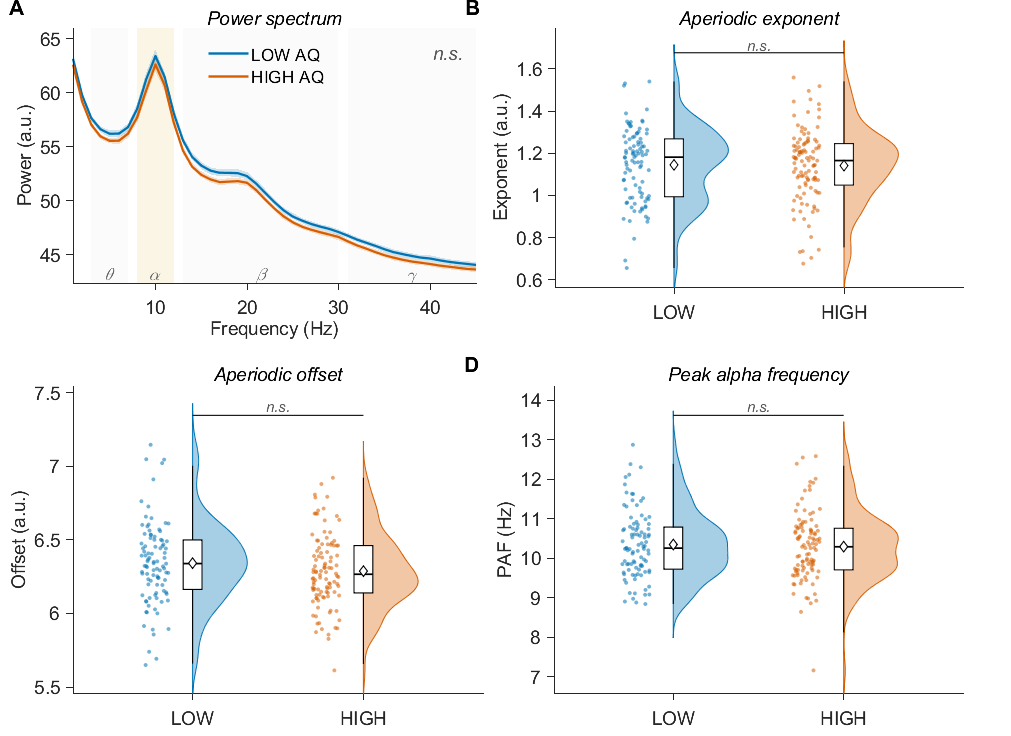

***Figure S2. Spectral control analyses in the occipito-central network: no group differences in oscillatory amplitude, aperiodic signal properties, or individual alpha frequency.***

*Same control pipeline as Figure 4, applied to the occipito-central set as a parallel analysis. (A) Power spectrum averaged across all occipito-central lines, with low-AQ (blue) and high-AQ (orange) participants shown as group means ± 1 SEM. (B–D) FOOOF-derived spectral parameters [aperiodic exponent (B), aperiodic offset (C), and peak alpha frequency (D)] aggregated per subject across lines and hemispheres, displayed as raincloud plots (half-violin kernel density, jittered individual points, compact boxplot, white diamond = group mean).None of the four metrics differed between AQ groups, with all Group and Group × Hemisphere terms yielding p > .10. This null pattern mirrors the one observed in the parieto-frontal network (Figure 4), indicating that AQ groups share comparable basic oscillatory profiles [power, 1/f structure, and alpha peak frequency] across both networks. The dissociation between the two networks reported in the main analyses therefore concerns the directional organization of traveling waves specifically, rather than any elementary spectral property of the underlying signal.*
